# Beta-hairpin Mechanism of Autoinhibition and Activation in the Kinesin-2 Family

**DOI:** 10.1101/2024.10.14.618219

**Authors:** Stephanie Webb, Katerina Toropova, Aakash G. Mukhopadhyay, Stephanie D. Nofal, Anthony J. Roberts

**Affiliations:** Sir William Dunn School of Pathology, University of Oxford, Oxford, UK; Institute of Structural and Molecular Biology, Department of Biological Sciences, Birkbeck, University of London, London, UK

**Author notes:** These authors contributed equally (Stephanie Webb, Katerina Toropova).

## Abstract

Members of the kinesin-2 family coordinate with other motors to power diverse physiological processes, but the structural mechanisms regulating kinesin-2 activity have been unknown. Distinctively, kinesin-2s canonically function as heterotrimers of two different motor subunits (e.g. Kif3A and Kif3B) and Kap3, but the role of heterotrimerization has yet to fully emerge. Here, we combine structural, cell biological, and single-molecule approaches to dissect kinesin-2 regulation as a heterodimer, heterotrimer, and quaternary complex with a cargo adaptor (APC). We identify a conserved motif in the tail of kinesin-2s (the beta-hairpin motif) that controls kinesin-2 motility by sequestering the motor domains away from their microtubule track, using striking molecular mimicry. Our data reveal how Kap3 binds via a multipartite interface with Kif3A and Kif3B, and – rather than activating motility directly – provides a platform on which cargo adaptors can engage and occlude the beta-hairpin motif. Together, these data articulate a structural framework for kinesin-2 activation, coordination with dynein, and adaptation for different biological functions.

## INTRODUCTION

Understanding how motor proteins coordinate to spatially organize the cytoplasm is a frontier in structural cell biology. The kinesin-2 family, which originated prior to the last eukaryotic common ancestor^1,2^, lies at the heart of this phenomenon, as its members work in a coordinated-fashion to transport cargo in diverse physiological processes^3,4^, including mRNA localization^5,6^, vesicular trafficking^7–9^, and intraflagellar transport (IFT)^10–14^. Loss-of-function in mammalian kinesin-2 subunits are embryonically lethal, often with situs inversus^15^, while recent studies identified patient mutations associated with ciliopathies^16,17^. Despite progress in understanding properties of kinesin-2 motor domains^18–22^, the structural mechanisms of kinesin-2 coordination and motility regulation are poorly understood.

Distinctively among kinesins, members of the kinesin-2 family typically function as heterotrimers^23^, in which two different motor subunits (Kif3A and Kif3B/Kif3C in humans) heterodimerize and associate with a third subunit, Kap3, which is built around Armadillo (ARM) repeats^24–27^. In mammals, Kif3AB-Kap3 functions in IFT and cytoplasmic transport^3,28^, while Kif3AC-Kap3 is implicated in neuronal transport^29^. Each motor subunit contains a kinesin motor domain that binds microtubules and hydrolyzes ATP^30,31^, coiled coil segments that mediate heterodimerization^32–34^, and a putatively disordered C-terminal tail^25,26^. The binding site of Kap3 is controversial, with one model positing that Kap3 binds to the coiled coil domain^35,36^ and others suggesting that binding occurs via the disordered tail of one or both motor subunits^25,26,29,37^. While essential for Kif3 function *in vivo*^13,38,39^, the precise role of Kap3 is also unclear. Reconstitution studies of *C. reinhardtii* kinesin-2 indicate that the motor subunits can move along microtubules without Kap3 but their velocity is increased in its presence^20^, whereas analysis of mouse proteins suggest that the kinesin-2 heterotrimer (Kif3AB-Kap3) is strongly autoinhibited^6^. Thus, the molecular basis and purpose of kinesin-2 heterotrimerization is yet to fully emerge.

Coordinated motor activity is particularly important in the elongated processes of neurons and within the confines of cilia and flagella^1^. In cilia, kinesin-2 powers anterograde movement of IFT trains and cargoes needed for cilia assembly to the tip, before dynein-2 drives return transport to the cell body^40,41^. To coordinate with other motors, members of the kinesin-2 family are thought to convert between autoinhibited and active states, with current models evoking folding of the molecule about a putative hinge in the coiled coil^3,18,20^. The molecular mechanism of motor autoinhibition is unclear, complicated by the question of whether one or both kinesin-2 motor subunits orchestrate the process. The issue is compounded by the fact that no autoinhibitory motif has been identified in the kinesin-2 subunits and the inhibitory sequence that regulates conventional kinesin-1 (the IAK motif^42^) is absent, suggesting that the regulatory mechanism may be novel. Valuable insights into kinesin-2 regulation have come from studies of homodimeric members of the family, Kif17 in humans and OSM-3 in *C. elegans*, which demonstrated that tail truncations, mutations at the putative hinge site, surface attachment, or binding of an associated factor (DYF-1/IFT70 in *C. elegans*) can stimulate motility^13,43–47^. Whether there is a unifying mechanism of regulation in the kinesin-2 family is unknown.

Here, we use cross-species structural analysis to identify a conserved motif in the tail of kinesin-2s (the β-hairpin motif), which was undetectable at the sequence level. We reveal that the β-hairpin motif inhibits kinesin-2 using a striking piece of molecular mimicry, in which negatively charged residues at the hairpin apex and adjacent coiled coil sequester the motor domains away from their microtubule track. Our data reveal how Kif3 exploits heterotrimerization: Kap3 binds via a multipartite interface with Kif3A and Kif3B and creates a surface where cargo adaptors can engage and occlude the β-hairpin motif. Together, these data provide a structural basis for kinesin-2 activation, coordination with dynein motors, and adaptation of family members for varied functions.

## RESULTS

### A β-hairpin motif predicted in the tail of kinesin-2 across eukaryotes

To investigate the basis for kinesin-2 autoregulation, we generated AlphaFold2 and AlphaFold3 (AF3) models for a range of kinesin-2 sequences from across eukaryotic supergroups^48^ (TSAR, Haptista, Cryptista, Archaeplastida, Amorphea, and Excavates), including heteromeric motors (*H. sapiens* KifA/B, *C. reinhardtii* FLA8/FLA10, and *C. elegans* KLP20/KL11) and homodimeric motors (*H. sapiens* Kif17 and *C. elegans* OSM-3). Notably, despite diversity in the length of the coiled coil segments and disordered tail region, a high-confidence feature common to all of the kinesin-2 models was present at the C-terminal end of the coiled coil, consisting of a short α-helix and a β-hairpin (hereafter referred to as ‘β-hairpin motif’) (**Figure 1A, Figure S1**). Structure-based alignment revealed that the apex of the β-hairpin invariantly bears negatively charged amino acids (**Figure 1B**). Sequence constraints for residues flanking the apex are looser, consistent with the β-hairpin structure, and indicating why the motif was not detected at the sequence level. However, there is strong conservation for aromatic residues in the +4 and –3 positions relative to the apex, which pack against each other (**Figure 1C**). The invariance of the negatively charged apex across eukaryotic supergroups, in both heteromeric and homodimeric kinesin-2s (**Figure 1C,D**), suggests it is ancestral and led us to hypothesize that it plays an important role in kinesin-2 mechanism.

**Figure 1.**
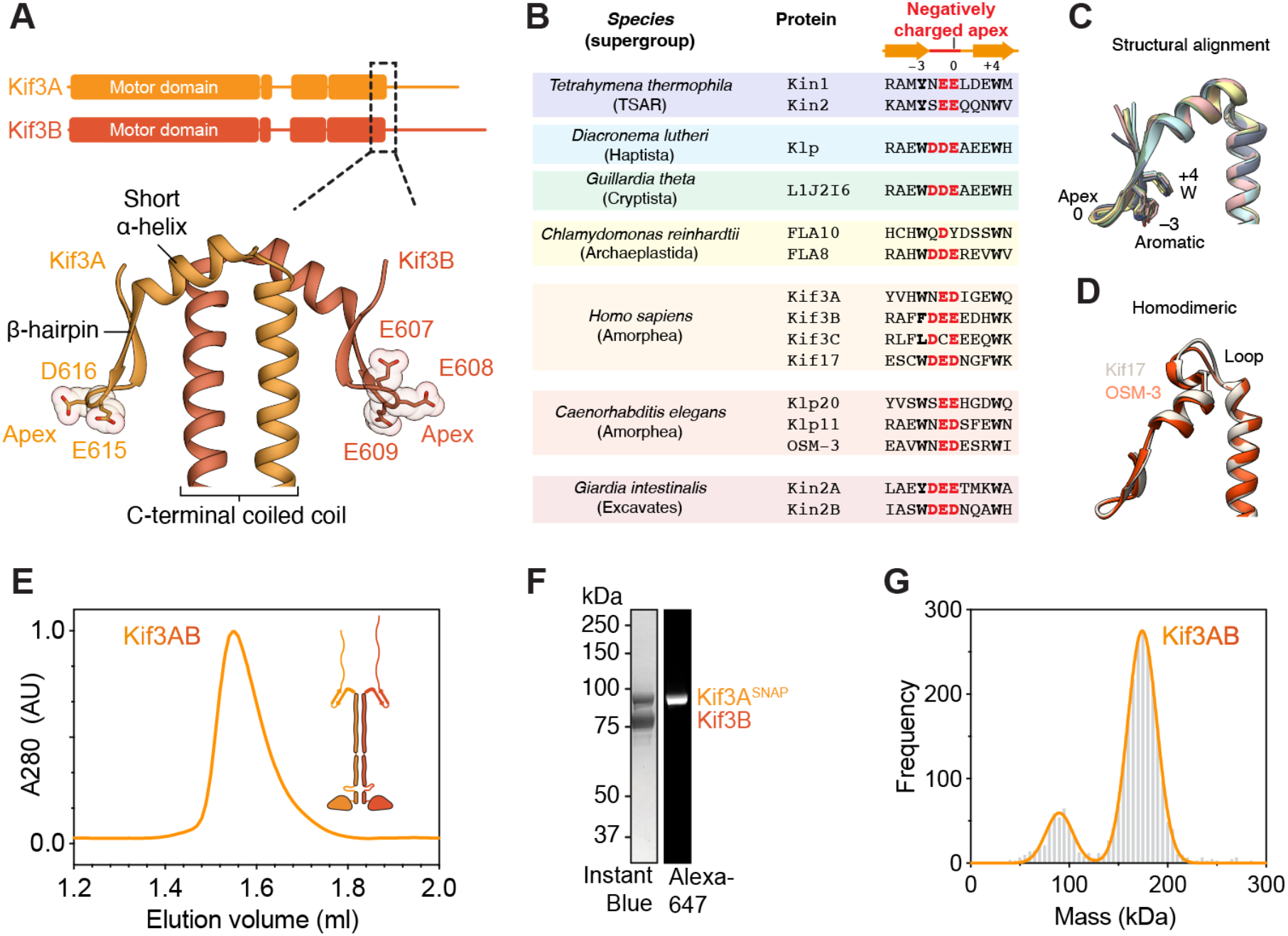
A β-hairpin motif predicted in the tail of kinesin-2 across eukaryotes. (A) Sequence diagrams of human kinesin-2 Kif3A and Kif3B. Beneath, AF3 model of the C-terminal coiled coil region and β-hairpin motif, with negatively charged residues at its apex shown in stick representation. (B) Structure-based sequence alignment of kinesin-2s from different eukaryotic supergroups. Negatively charged residues at the apex of the β-hairpin are in red. Conserved aromatic residues in the –3 and +4 positions relative to the apex are in bold. (C) Aligned AF3 models of the β-hairpin in different kinesin-2s, colored as in panel B. (D) AF3 models of homodimeric kinesin-2 Kif17 and OSM-3, which both feature an additional loop between the coiled coil and short α-helix preceding the β-hairpin. (E) Size-exclusion chromatogram of reconstituted Kif3AB heterodimer with schematic of the construct inset. (F) SDS-PAGE of peak size-exclusion chromatography fraction after labeling SNAP-tagged Kif3A with Alexa 647 fluorophore. (G) Mass photometry of purified Kif3AB. Main peak (174 ± 15 kDa, mean ± SD) is consistent with Kif3AB heterodimer mass (166 kDa).

### Disruption of the β-hairpin motif activates kinesin-2 motility

To investigate the role of the β-hairpin motif, we co-expressed and purified *H. sapiens* Kif3AB from insect cells. A SNAP_f_ tag on Kif3A enabled covalent labeling with biotin (for surface immobilization in multi-motor microtubule gliding assays) or bright fluorophores (for single-molecule motility assays). Kif3AB was separated from excess ligand using size-exclusion chromatography (**Figure 1E**), yielding purified (**Figure 1F**) heterodimeric (**Figure 1G**) protein.

In single-molecule motility assays, full-length Kif3AB did not detectably bind to or move along microtubules in the presence of ATP, suggestive of strong autoinhibition (**Figure 2A**, upper panel). In the presence of AMPPNP, which traps the kinesin motor domain in a high microtubule affinity state^49^, full-length Kif3AB bound statically to microtubules (**Figure 2A**, lower panel). To test if the lack of motility and microtubule binding in ATP conditions is due to autoinhibition at the single-molecule level, we biotinylated full-length Kif3AB and attached multiple motors to a neutravidin-coated surface via their tail (**Figure 2B**). Upon addition of microtubules and ATP, surface-immobilized Kif3AB molecules powered robust microtubule gliding at 444 ± 26 nm s^-1^ (mean ± SEM from 3 separate experiments) (**Table 1**). These data indicate that full-length human Kif3AB is capable of driving motility when multiple motors are anchored via their tail but is strongly autoinhibited as a single molecule in solution.

**Table 1.** Values show mean ± SEM from 3 separate experiments. More than 100 motile events were analyzed per experiment. No motile events were observed for Kif3A-Kif3B, Kif3A^ΔC38^-Kif3B, or Kif3A^ΔC59^-Kif3B, suggestive of strong autoinhibition. Landing rates are not given for Kif3A^KK^-Kif3B, Kif3A-Kif3B^KKK^, or Kif3A^KK^-Kif3B^KKK^ as accurate protein concentrations not measurable (low protein yield). Landing rate for these constructs is broadly comparable to Kif3A-Kif3B-Kap3-APC.

| **Construct** | **Velocity**<br>**(nm s<sup>-1</sup>)** | **Landing rate**<br>**(μm<sup>-1</sup> nM<sup>-1</sup> min<sup>-1</sup>)** | **Run length**<br>**(μm)** |
| --- | --- | --- | --- |
| Kif3A-Kif3B | — | — | — |
| Kif3A<sup>ΔC38</sup>-Kif3B | — | — | — |
| Kif3A<sup>ΔC59</sup>-Kif3B | — | — | — |
| Kif3A<sup>ΔC89</sup>-Kif3B | 945 ± 30 | 0.55 ± 0.04 | 0.27 ± 0.01 |
| Kif3A<sup>ΔC103</sup>-Kif3B | 933 ± 23 | 0.60 ± 0.07 | 0.27 ± 0.02 |
| Kif3A<sup>ΔC103</sup>-Kif3B<sup>ΔC39</sup> | 990 ± 4 | 0.56 ± 0.06 | 0.28 ± 0.03 |
| Kif3A<sup>ΔC103</sup>-Kif3B<sup>ΔC90</sup> | 939 ± 4 | 3.30 ± 0.20 | 0.20 ± 0.02 |
| Kif3A<sup>ΔC103</sup>-Kif3B<sup>ΔC114</sup> | 950 ± 15 | 3.40 ± 0.26 | 0.21 ± 0.01 |
| Kif3A<sup>ΔC103</sup>-Kif3B<sup>ΔC156</sup> | 903 ± 30 | 2.70 ± 0.18 | 0.33 ± 0.09 |
| Kif3A<sup>KK</sup>-Kif3B | 730 ± 71 | — | 0.55 ± 0.04 |
| Kif3A-Kif3B<sup>KKK</sup> | 740 ± 75 | — | 0.64 ± 0.04 |
| Kif3A<sup>KK</sup>-Kif3B<sup>KKK</sup> | 693 ± 82 | — | 0.60 ± 0.06 |
| Kif3A-Kif3B-Kap3-APC | 792 ± 23 | 0.09 ± 0.01 | 0.78 ± 0.02 |

**Figure 2.**
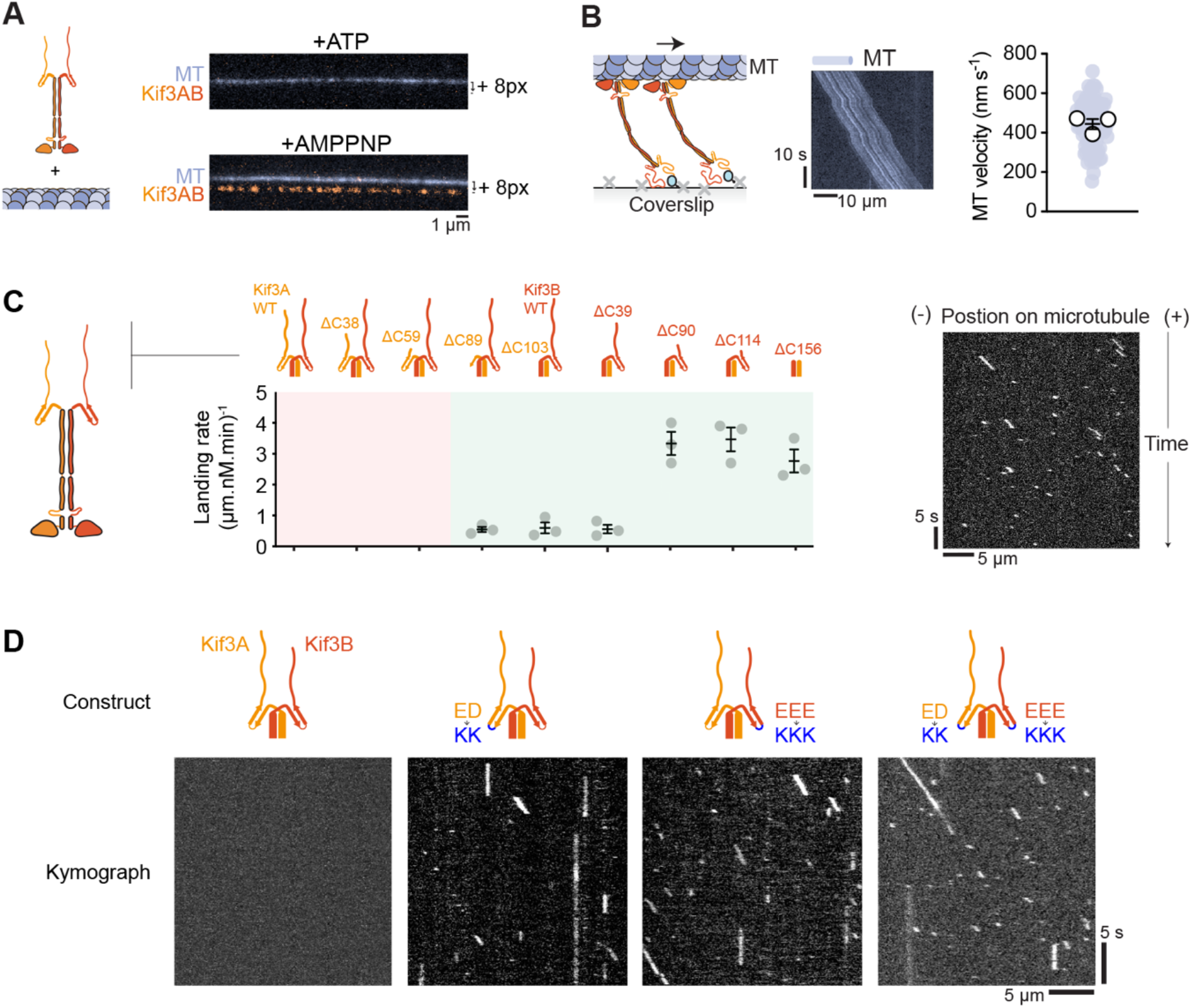
The β-hairpin motif mediates kinesin-2 autoinhibition. (A) Composite TIRF images of 488 nm channel (MT) and 640 nm channel (Alexa-647-labeled Kif3A) offset in y-axis by 8 pixels. Top, with 1 mM ATP. Bottom, with 1 mM AMPPNP. MT; microtubule. (B) Left, schematic of MT gliding assay with biotinylated Kif3AB. Middle, example kymograph. Right, plot of gliding velocities from three technical replicates. Colored circles; individual data points. White circles; average from each separate experiment. Lines; mean (±SEM), n = 86 microtubules. (C) Left, microtubule landing rate of Alexa-647-labeled Kif3AB on microtubules. Constructs were sequentially truncated from their C-termini as indicated in schematics above plots. Measurements were taken from three technical replicates, n > 100 motile events analyzed per experiment. Gray circles; average from each experiment. Lines; mean (±SEM). Right, example kymograph of Kif3A^ΔC103^B^ΔC156^. (D) Example kymographs of Kif3AB β-hairpin mutants. Motility parameters for all constructs are given in Table 1.

To home in on the region/s of Kif3AB responsible for autoinhibition, we made a series of C-terminal truncations (which did not affect heterodimerization; **Figure S2A)**. Truncation of 38 or 59 putatively disordered amino acids from the C-terminus of Kif3A had no effect on Kif3AB single-molecule motility, which remained strongly autoinhibited (**Figure 2C**). Strikingly, however, truncations within or beyond the β-hairpin motif of Kif3A (89 and 103 amino acids, respectively), activated the motor, which bound to and moved along microtubules with an average velocity of ∼940 nm s^-1^, landing rate of ∼0.6 μm^-1^ nM^-1^ min^-1^, and run length of ∼0.3 μm (**Table 1**). These data indicate that disruption or removal of the Kif3A β-hairpin is sufficient to release Kif3AB autoinhibition. We next examined if C-terminal truncations of Kif3B could further stimulate motility in constructs lacking the Kif3A β-hairpin (**Figure 2C**). While truncation of 39 amino acids from the C-terminus of Kif3B had no effect, removal of 90 amino acids increased the landing rate to ∼3.2, while velocity and run length remained unchanged (**Table 1**). Larger Kif3B C-terminal truncations of 114 or 156 amino acids (the latter removing the Kif3B β-hairpin motif) did not elicit further increases in motility. These results indicate that a segment of the Kif3B tail can suppress landing rate while raising the question of whether disrupting one β-hairpin motif in Kif3AB is sufficient for activation.

To directly probe the contributions of the Kif3A and Kif3B β-hairpin motifs to autoinhibition, we mutated the conserved negatively charged amino acids at their apex to lysine in Kif3A and Kif3B individually or in tandem (**Figure 2D**). Mutation of the Kif3A β-hairpin activated Kif3AB motility, as did mutation of the Kif3B β-hairpin, yielding motors that moved with similar yet slightly longer run lengths (∼0.6 μm) and lower velocity (∼730 nm s^-1^) compared to constructs activated by truncation. In agreement with the truncation experiments, there was no additive effect when both Kif3A and Kif3B β-hairpins were mutated (**Figure 2D, Table 1**). Based on these data, we conclude that 1) the β-hairpin motif has a fundamental role in Kif3AB autoinhibition and 2) disruption of one β-hairpin, from either Kif3A or Kif3B, is sufficient to destabilize the autoinhibited conformation of Kif3AB.

### Impact of disrupting the β-hairpin on Kif3 cellular function

We next examined the role of the β-hairpin motif in Kif3 cellular activity, using the function of Kif3 in ciliary transport in IMCD-3 cells, a mouse kidney cell line, as an exemplar. IMCD-3 cells form a primary cilium upon serum starvation (cilia length of 3.3 ± 0.2 μm [mean ± SEM]) (**Figure 3A**), in which Kif3 powers outward movement of IFT trains and cargoes to the tip and is then transported back to the cell body on dynein-2 powered trains^50^ (or by diffusion in *C. reinhardtii*^51^). We used CRISPR/Cas9 to generate a knockout (KO) of Kif3B (**Figure S2B**), which ablated cilia formation (**Figure 3A, S2C**). Stable expression of wild-type Kif3B-mScarlet in the KO cells rescued to the cilia length to 3.02 ± 0.02 μm, confirming that the loss of cilia is due to Kif3B deletion. Live-cell total internal reflection fluorescence (TIRF) microscopy revealed that Kif3B-mScarlet signal localized approximately evenly along the length of cilia (**Figure 3B**, upper panel), as quantified in lines scans of the time-averaged fluorescence intensity (**Figure 3B**, lower panel). A Kif3B construct unable to bind to IFT trains and impaired in Kap3 binding^37^ (Kif3B^ΔC156^) failed to restore cilia length, as expected (**Figure 3A**), despite being expressed at comparable levels to the wild-type construct (**Figure S2D**). The Kif3B β-hairpin mutant rescued cilia length (**Figure 3A**), suggesting it is capable of delivering building blocks to the tip of the cilium to support ciliogenesis. However, unlike the wild-type construct, the Kif3B β-hairpin mutant accumulated in bright puncta at the ciliary tip (**Figure 3B**). These results indicate that the constitutively active Kif3B β-hairpin mutant can power transport to the tip of cilia, to support ciliogenesis, but is not effectively returned to the cell body, highlighting autoinhibition via the β-hairpin motif as crucial for Kif3 spatial regulation and coordination with dynein-2.

**Figure 3.**
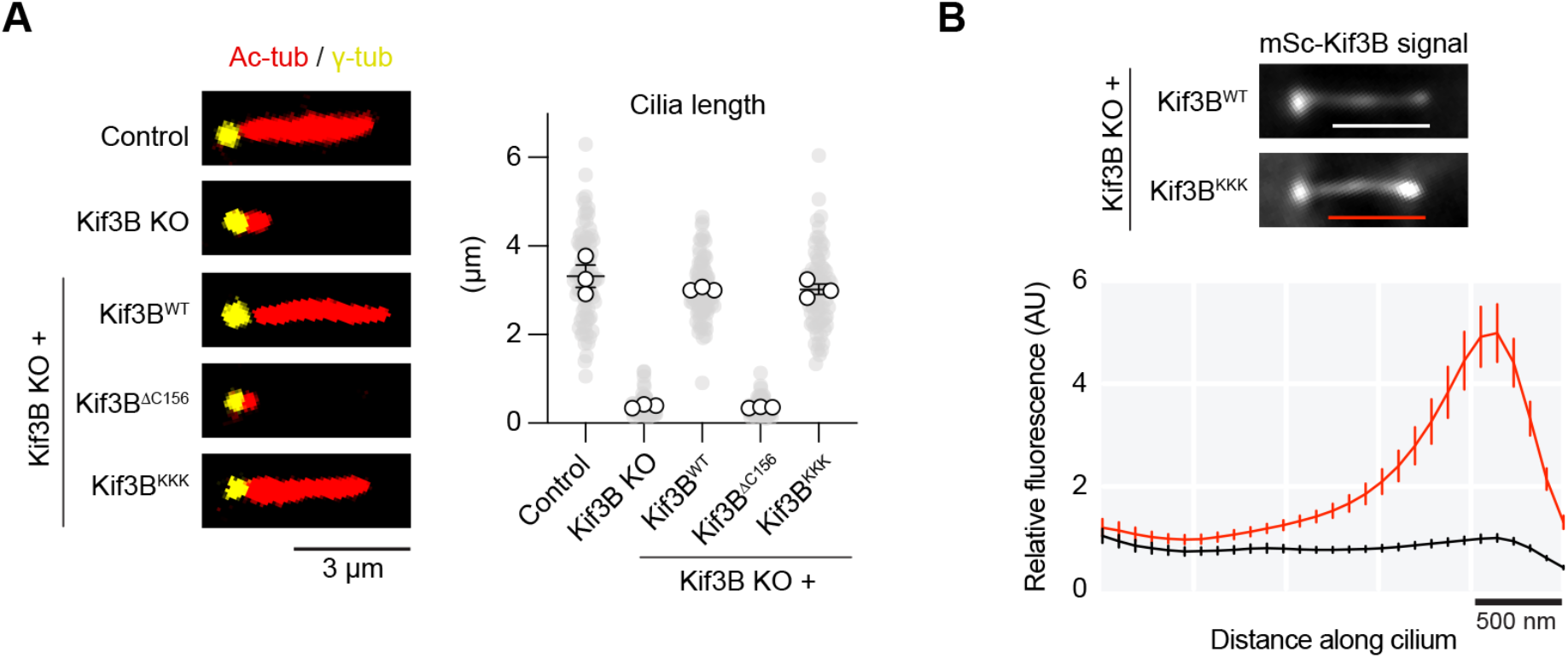
The β-hairpin motif is crucial for Kif3 recycling and spatial regulation in intraflagellar transport. (A) Left, representative IMCD-3 cilia in control cells and Kif3B KO cells stably expressing indicated constructs. Cells were immunofluorescently labeled for acetylated tubulin (red) marking the ciliary axoneme and gamma tubulin (yellow) marking the basal body. Right, quantification of cilia length from three separate experiments. Gray circles; individual data points. White circles; average from each separate experiment. Lines; mean (±SEM). Control n = 86; Kif3B KO n = 76; Kif3B^WT^ n = 78, Kif3B^ΔC156^ n = 76, Kif3B^KKK^ n = 74 cilia analyzed. One-way ANOVA followed by Kruskal-Wallis test values, control vs. Kif3B KO p < 0.0001; control vs. Kif3B^WT^ p > 0.1; control vs. Kif3B^ΔC156^ p < 0.0001; control vs. Kif3B^KKK^ p > 0.1. (B) Top, representative images of mScarlet (mSc) tagged Kif3B^WT^ and Kif3B^KKK^ constructs expressed in Kif3B KO cells. Bottom, plot of average mSc-Kif3B fluorescent signal from line scans along the cilium length (bars in top panel indicate distance analyzed), aligned at the ciliary tip (peak in the Kif3B^WT^ trace). Values are normalized relative to Kif3B^WT^ peak value. Traces show mean intensity ± SEM; n = 42 (Kif3B^WT^), 51 (Kif3B^KKK^) cilia measured from three separate experiments.

### Architecture of the Kif3 heterotrimer

We next sought to understand how Kif3AB binds to Kap3 to form the Kif3 heterotrimer. We purified human Kap3 from insect cells and reconstituted the heterotrimeric complex with full-length Kif3AB, as verified by size-exclusion chromatography (**Figure 4A,B**) and mass photometry (**Figure S3A**). In single-molecule motility TIRF assays, Kif3AB-Kap3 did not bind to or move along microtubules in the presence of ATP, suggesting that Kap3 binds Kif3AB in a way that allows autoinhibition to persist (**Figure 4C, upper panel**). In the presence of AMPPNP, Kif3AB and Kap3 bound statically to microtubules and co-localized with each other (**Figure 4C, lower panel**), showing that the complex is stable at the nanomolar concentrations of the TIRF assay.

**Figure 4.**
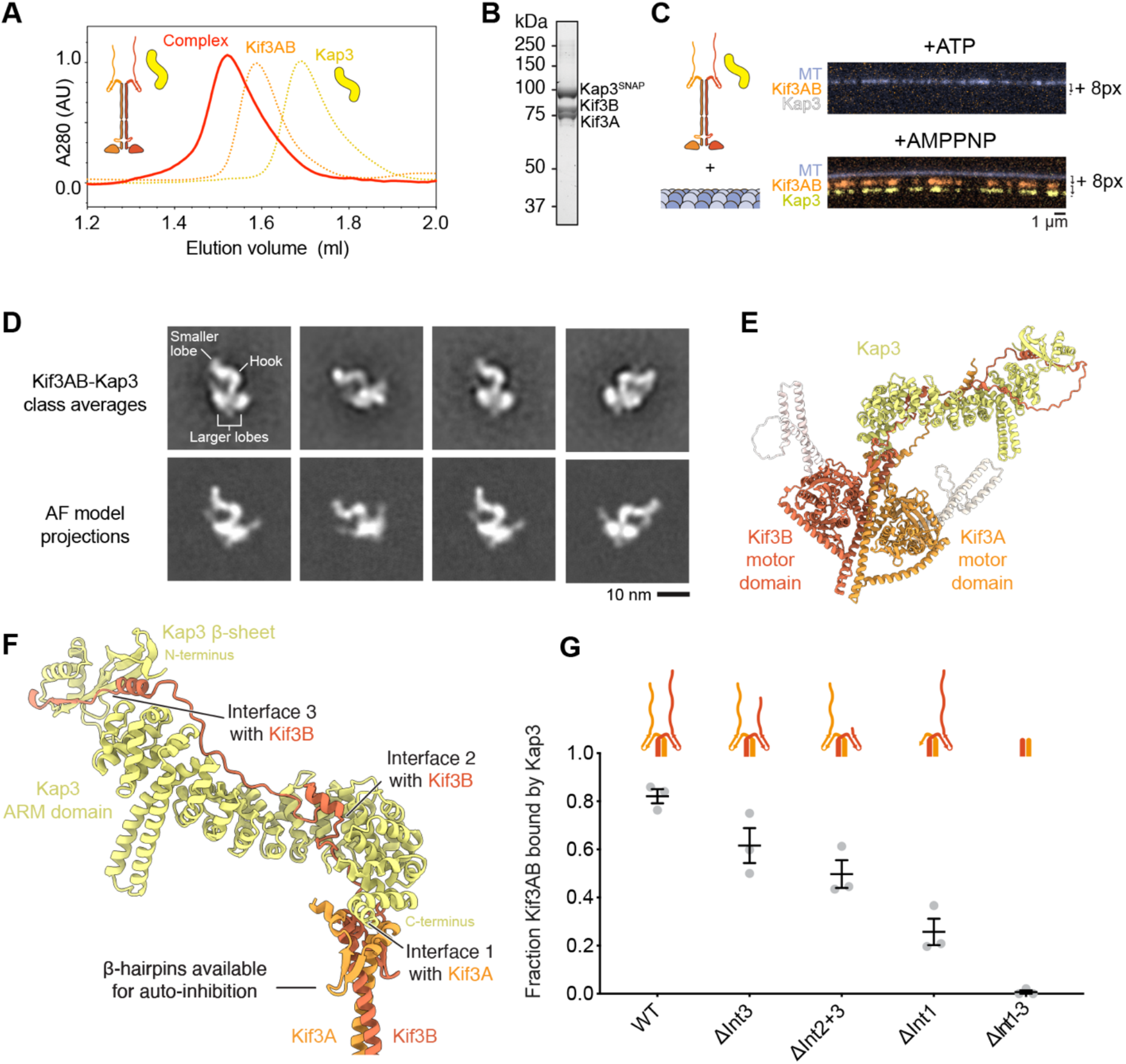
Kap3 makes a multipartite interaction with Kif3AB that permits autoinhibition. (A) Size-exclusion chromatogram of reconstituted Kif3AB–Kap3 complex in red with schematic alongside. Normalized Kif3AB and Kap3 traces are shown for comparison (dashed orange and yellow lines respectively). (B) SDS-PAGE of peak size-exclusion chromatography fraction. (C) Top, composite TIRF image of 488 nm channel (MT) and 640 nm channel (Alexa-647-labeled Kif3A) offset in y-axis by 8 pixels, assay with 1.5 nM Kif3AB, 7.5 nM Kap3 (unlabeled) and 1 mM ATP. The presence of Kap3 does not activate Kif3AB microtubule binding or motility. Bottom, as for above but with 561 nm channel (TMR-labeled-Kap3) offset in y-axis by 16-pixels, assay with 1 nM Kif3AB, 1 nM Kap3 and 1 mM AMPPNP. Kif3AB and Kap3 co-localize in a complex. (D) Top, example class averages of Kif3AB-Kap3 complex, with features labeled. Bottom, corresponding AF3 model projections. (E) Ribbon representation of Kif3AB-Kap3 AF3 model. F) Close-up of composite binding interface between Kap3 and Kif3AB C-terminal region. The three main interfaces are labeled. Kif3AB β-hairpins are not occluded by Kap3 binding. (G) Plot of the fraction of TMR-labeled Kap3 co-localizing with Alexa-647-labeled Kif3AB for each indicated construct: ΔInt3 (Kif3A-Kif3B^ΔC90^), ΔInt2+3 (Kif3A-Kif3B^ΔC114^), ΔInt1 (Kif3A^ΔC89^-Kif3B), ΔInt1-3 (Kif3A^ΔC103^-Kif3B^ΔC156^). Measurements were taken from three technical replicates. Gray circles; average from each separate experiment. Lines; mean (±SEM), n > 100 molecules analyzed per construct.

To gain insight into how Kap3 binds to Kif3AB in a manner that allows autoregulation, we examined the structure of the heterotrimer using single-particle negative stain electron microscopy (EM) (**Figure 4D, Figure S3B**). Kif3AB-Kap3 displayed a compact shape comprising a hook-like density, matching the expected dimensions of the Kap3 ARM repeat α-solenoid^52^, with a small lobe at one end and two larger lobes at the other. The larger lobes are consistent with the size of the kinesin motor domains, and a fine structure is occasionally visible projecting out from them. To help interpret these features and generate an atomic hypothesis for the binding mode, we made an AlphaFold3 model of the Kif3AB-Kap3 heterotrimer (**Figure 4E**). Projections of this model showed striking concordance with the experimental EM class averages (**Figure 4D**) and support assignment of the hook-like feature as Kap3 and the two larger lobes as the Kif3A and Kif3B motor domains (**Figure 4E**).

In this high-confidence model, Kap3 binds to Kif3AB via multiple interfaces (**Figure 4F, Figure S3C**). First, the C-terminal end of Kap3’s ARM domain interacts with a pocket formed by short α-helices of Kif3A flanking the β-hairpin motif (Interface 1). Notably, this contact leaves the apex of the β-hairpin motifs free to interact with the Kif3AB motor domains (described below), indicating why the Kif3 heterotrimer remains autoinhibited. Second, the disordered Kif3B tail runs along the inner surface of the Kap3 ARM α-solenoid (Interface 2). Notably, this site encompasses a conserved phosphorylation site in Kif3B^53^ (see Discussion). Third, the C-terminal region of the Kif3B tail forms a β-strand and α-helix that interact with a 3-stranded β-sheet at the Kap3 N-terminus (Interface 3). This β-sheet corresponds to the smaller lobe at the distal end of Kap3 observed in our EM class averages. Finally, the disordered Kif3A tail also interacts with the Kap3 ARM domain, although much less extensively than the Kif3B tail (**Figure S3C**).

To test the importance of these interfaces in Kap3 binding, we performed a sensitive single-molecule binding assay, in which Kif3AB and Kap3 were labeled with different color fluorophores (**Figure 4C,G**). Whereas Kap3 co-localized with 82 ± 3% of full-length Kif3AB molecules (mean ± SEM), ablation of Interface 3 by truncation of Kif3B tail reduced co-localization to 62 ± 7%, which was further reduced to 50 ± 6% by removal of Interface 2. Removal of Interface 1, by truncation of the Kif3A, had a stronger effect, reducing co-localization to 26 ± 6%. Finally, removal of all the predicted interfaces, while retaining the N-terminal portion of Kif3AB, abolished co-localization, showing that Kap3 does not bind appreciably to the Kif3AB coiled coil or motor domains. We conclude that Kap3 makes a multipartite interaction with the C-terminal regions of both Kif3A (Interface 1) and Kif3B (Interfaces 2 and 3).

### Structural basis for Kif3 autoinhibition

Our EM class averages, mutagenesis data, and AlphaFold analysis suggest a structural framework for Kif3 autoinhibition (**Figure 5**). In this model, the Kif3A and Kif3B motor domains lie either side of the C-terminal coiled coil segment, interacting with 1) their respective β-hairpin motif and 2) the C-terminal Kif3AB coiled coil (**Figure 5A**). Both interactions have a major electrostatic component and mimic the charged-based interactions that the kinesin motor domain makes with α/β-tubulin. A basic patch in the Kif3A motor domain (involving Lys301 and Arg304) interacts with the acidic apex of the Kif3A β-hairpin (Glu612, Asp613), echoing the interaction with β-tubulin^54^ (**Figure 5B**). A second basic patch in the Kif3A motor domain (involving Lys50, Lys59, Arg256, and Arg337) interacts with a cluster of electronegative residues in the C-terminal coiled coil (Kif3A Glu559, Asp562, Glu566, and Glu570) mimicking the interaction with α-tubulin^54^ (**Figure 5B**). Similar, but non-identical interactions are made by the Kif3B motor domain (**Figure 5C**), suggesting why chimeras in which the Kif3A motor domain is transplanted onto the Kif3B C-terminal region, and *vice versa*, are not autoinhibited^17,18^. Together, these interactions hold the motor domains of Kif3A and Kif3B in a pseudo 2-fold symmetrical arrangement with their microtubule-binding domains occluded, consistent with the lack of Kif3AB microtubule binding observed in our motility assays (**Figure 5D**). Our data suggest that this autoinhibited conformation is meta-stable, as AMPPNP or mutagenesis of only one β-hairpin is sufficient to disfavor it (**Figure 2**), indicating that it could be a target for regulation.

**Figure 5.**
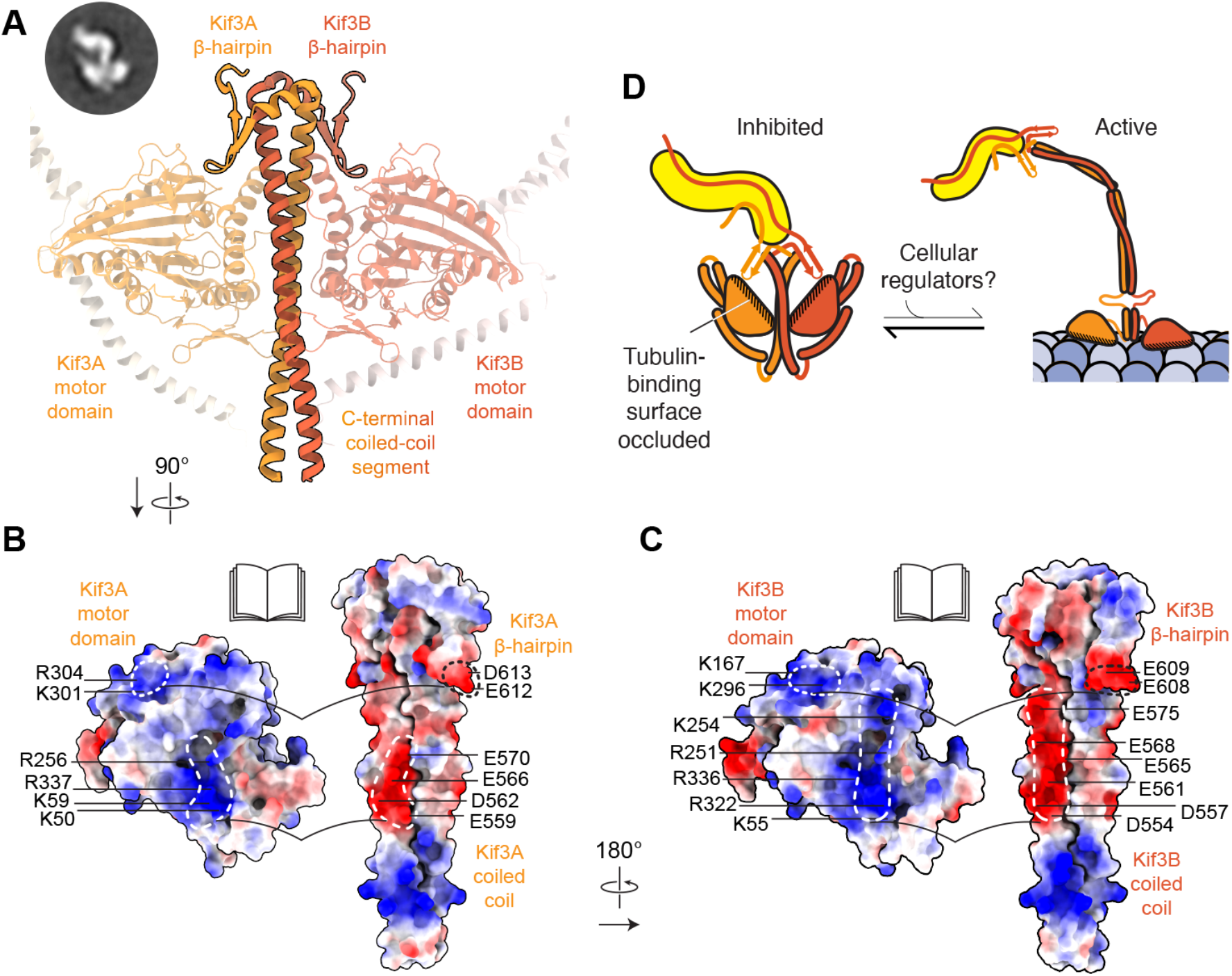
Structural basis for Kif3 autoinhibition. (A) Close-up of AF3 model of autoinhibitory Kif3AB interface in cartoon representation and colored as for previous figures. Key features labeled. Negative stain EM class average from Figure 4D shown inset. (B) Open-book representation of binding interface between Kif3A motor domain and Kif3A β-hairpin and coiled coil. The model is shown in surface representation and colored by electrostatic potential with key residues annotated. Note the complementary charges between motor domain (positively charged; blue) and β-hairpin and coiled coil (negatively charged; red). (C) Open-book representation of binding interface between Kif3B motor domain and Kif3B β-hairpin and coiled coil. (D) Schematic model for kinesin-2 autoinhibition. Kif3A and B motor domains fold back to interact with β-hairpin motifs occluding the tubulin binding interface of each motor. Kap3 binds C-terminal regions of Kif3AB and is not involved in autoinhibition, remaining available to bind other factors that may activate kinesin-2.

### Activation of Kif3 by adaptor binding to the β-hairpin motif

How might the subunits of Kif3 be acted on by cellular factors to elicit activation? To address this question, we purified Adenomatous Polyposis Coli (APC), a well-characterized adaptor that couples Kif3 to mRNA cargoes in neurons and whose ARM domain (APC^ARM^) is involved in Kif3 motility regulation^6,29,52,55^. Size-exclusion chromatography demonstrated that APC^ARM^ binds stably to the Kif3AB-Kap3 heterotrimer (**Figure 6A**), but weakly to Kif3AB alone (**Figure S4A**), suggesting that Kap3 forms an important part of the Kif3-APC^ARM^ interface. Notably, single-molecule TIRF assays showed that APC^ARM^ was sufficient to activate the motility of Kif3AB-Kap3, which bound to and moved along microtubules (**Figure 6B**) with motile parameters akin to constructs activated by β-hairpin mutation (**Table 1**). In contrast, APC^ARM^ did not activate the motility of Kif3AB alone, consistent with the binding assay (**Figure S4A**,**B**). Together, these data indicate that APC^ARM^ activates Kif3 in a Kap3-dependent manner.

**Figure 6.**
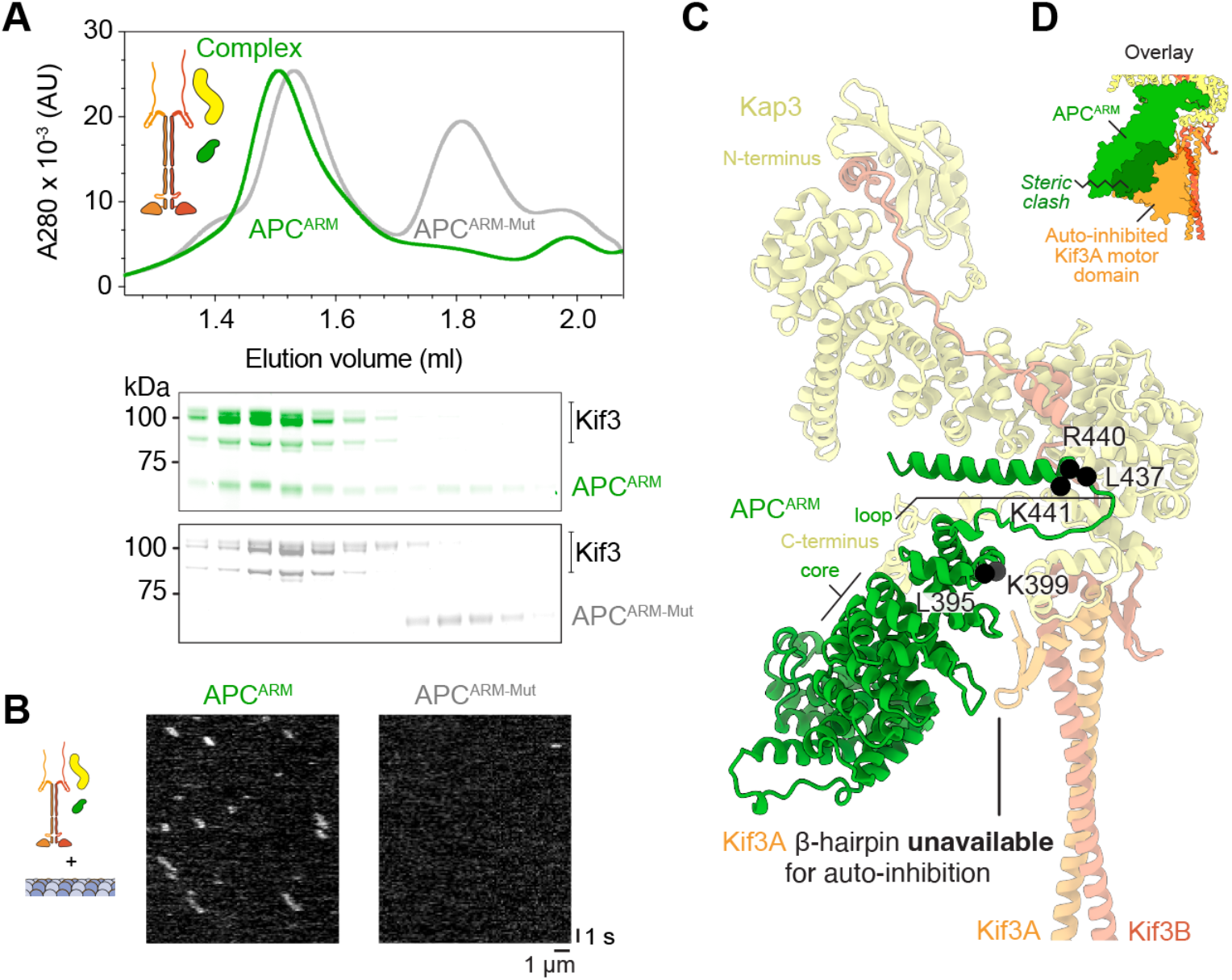
Kap3-dependent activation of Kif3 by adaptor binding to the β-hairpin motif. (A) Top, size-exclusion chromatogram of reconstituted Kif3AB–Kap3 complex in presence of either APC^ARM^ (green) or APC^ARM-Mut^ (grey). Bottom, SDS-PAGE of size-exclusion chromatography fractions. (B) Example kymographs of Kif3AB-Kap (Alexa-647-labeled Kif3A) in presence of microtubules and APC^ARM^ or APC^ARM-Mut^. (C) AF3 model of APC^ARM^ in complex with Kif3AB-Kap3. Conserved APC^ARM^ residues that interact with the complex via Kap3 are shown as black spheres and annotated (mutated in APC^ARM-Mut^). APC^ARM^ occludes the Kif3A β-hairpin. (D) Equivalent view to (C) with autoinhibited Kif3A motor domain binding site overlaid, showing steric clash with APC^ARM^. APC^ARM^ and Kif3A motor domain shown in surface representation.

To explore the basis for Kif3 activation by APC^ARM^, we generated an AlphaFold3 model of their interface (**Figure 6C**). In this model, APC^ARM^ binds a composite surface consisting of the C-terminal region of the Kap3 ARM domain and, strikingly, the β-hairpin motif of Kif3A, with each interaction supported by low predicted aligned error (**Figure S4C**). The interaction with the Kap3 ARM domain involves the core of APC^ARM^ and a loop that protrudes from its C-terminal end. The disordered C-terminal region of Kap3 also contacts the internal surface of APC^ARM^. Notably, the binding footprint of APC^ARM^ sterically clashes with the position of the Kif3A motor domain in the autoinhibited conformation of Kif3 and occludes the Kif3A β-hairpin (**Figure 6C**,**D**), suggesting why APC^ARM^ activates Kif3, as observed in our motility assays (**Figure 6B**). The notion that occlusion of one β-hairpin is sufficient for activation is consistent with our data showing Kif3AB autoinhibition can be relieved by mutagenesis of a single β-hairpin (**Figure 2D**). To test this activation model, we mutated conserved surface residues in APC^ARM^ at the putative interface with Kif3. Whereas wild-type APC^ARM^ robustly co-eluted with Kif3 in a size-exclusion chromatography assay, binding was effectively abolished by the interface mutations (**Figure 6A**). Moreover, in contrast to wild-type APC^ARM^ that activated Kif3 motility, the APC^ARM^ interface mutant completely failed to relieve Kif3 autoinhibition in single-molecule TIRF assays (**Figure 6B**), demonstrating that binding at this interface is the cause of Kif3 activation. Together, these results indicate that APC^ARM^ activates Kif3 motility in a Kap3-dependent manner by binding to a composite interface comprising Kap3 and the Kif3A β-hairpin.

## DISCUSSION

Here, using purified human proteins, EM, AlphaFold, single-molecule TIRF, and CRISPR-Cas9 genome editing, we have dissected the mechanism of Kif3 motility regulation and heterotrimerization. Our results have implications across the kinesin-2 family: one of the “toolbox” motors present in the last eukaryotic common ancestor, whose extant members function in IFT and cilia self-assembly, Hedgehog and Wnt signaling, vesicle transport, and axonal mRNA localization, among other vital physiological roles^1–3,56,58,59^.

We find that a conserved β-hairpin motif mediates autoinhibition in kinesin-2 using a remarkable piece of molecular mimicry, in which negatively charged residues at the hairpin apex and adjacent C-terminal coiled coil mimic the charge distribution of α/β-tubulin and restrict access of the motor domains to their microtubule track. This structural mechanism, derived from our EM data, pseudo-atomic model, and mutagenesis, is compatible with foundational images of sea urchin kinesin-2 in a compact morphology observed by rotary shadow EM^25^. The ubiquity of the β-hairpin motif we find predicted in kinesin-2s across the eukaryotic supergroups, and its absence in other kinesin families, suggests it may serve as a useful tool for identifying kinesin-2 members, for example in poorly annotated proteomes in which motor domain-based classification is ambiguous. One rare exception is trypanosomatids, the only group we have found thus far to lack a predicted β-hairpin in its kinesin-2 sequences (which also display other divergent features^57^), suggesting that the regulatory mechanism in these human parasites may be unique.

The β-hairpin mechanism we describe provides a basis for why a C-terminal Kif3A fragment (which we can now see includes the β-hairpin motif) inhibits Kif3AB *in trans*^60^, and why mutations at the putative hinge site in the coiled coil relieve kinesin-II autoinhibition^17,18,43,45^ (as they would disfavor access of the β-hairpins to the motor domains). However, our EM data and analysis of the coiled coil (**Figure S5**) suggest that formation of the inhibited state is more complex than folding of the molecule about the putative hinge. Rather, our data indicate that in the compact, inhibited state we observe, the coiled coil segments proximal to the motor domains separate from one another and pack against themselves to shield their hydrophobic seams, whereas formation of the active state would involve zippering of these α-helical segments together into a canonical coiled coil (**Figure S5**).

Our data indicate that heterotrimeric kinesin-2 intrinsically exists in a compact, β-hairpin-engaged state, but transiently samples the extended (β-hairpin-disengaged) state, consistent with sedimentation analysis^25^ and providing a plausible explanation for previous observations. For example, the finding that *C. reinhardtii* FLA10/FLA8 can move along microtubules whereas mouse and human Kif3AB are strongly autoinhibited can be interpreted by differing buffer conditions or phosphorylation affecting the equilibrium position between the states^6,20,61^. It is also of note that FLA10 has fewer negative charges at the apex of its β-hairpin compared to many of its Kif3A orthologs, which could favor the β-hairpin-disengaged state (**Figure 1B**). *In vivo*, we think it likely that the β-hairpin-engaged state predominates for isolated kinesin-2, because β-hairpin-mediated autoinhibition provides the opportunity for controlled activation of motility. In line with this proposal, Kif3AB and homodimeric kinesin-2, Kif17, exist in an autoinhibited state when expressed in cells^17,45^.

Our data suggest how kinesin-2 exploits heterotrimerization: Kap3 binds via a multipartite interface with Kif3A and Kif3B, and – rather than activating motility directly – provides a surface where cargo adaptor, exemplified by APC in this study, can stably engage and occlude the β-hairpin motif. APC activates kinesin-2 for mRNA transport in neurons^6^ and we anticipate that other adaptors activate kinesin-2 in different biological contexts using analogous mechanisms. For example, binding of heterotrimeric kinesin-2 to the IFT-B complex may contribute to the initiation of anterograde IFT^62^; in *C. elegans*, where homodimeric kinesin-2 OSM-3 drives anterograde transport along the distal ciliary segment^13,14,63^, binding of OSM-5(IFT88)/DYF-1(IFT70)/DYF-6(IFT46)/OSM-6(IFT52) to the extended C-terminal region of OSM-3 could occlude the β-hairpin motif and explain the observed motor activation^46^. It will be interesting to examine if distinct adaptors underlie Kif3-mediated transport of other cargoes, a striking example being the regulated transport of melanosomes that underlies the color changes of amphibians^9^.

We show that, rather than binding to the coiled coil or motor domains, Kap3 engages Kif3AB via multiple interfaces in their C-terminal regions. Among the interfaces between Kap3 and Kif3AB we describe, Interface-2 and -3 involving the Kif3B tail are particularly interesting from a regulatory perspective. Interface 2 includes a conserved phosphorylation site, which in C. *reinhardtii* influences IFT turnaround^53^. Interface 3 is notable because it is a main point of departure between Kif3AB-Kap3, which functions in IFT, and Kif3AC-Kap3, which does not^28^. We find that Kif3AC is predicted to interact with Kap3 using Interfaces 1–2, consistent with binding experiments^29^ (**Figure S3D**), but lacks Interface 3 (**Figure S3C**). Thus, Interface 3, which involves the C-terminal region of Kif3B contributing a β-strand and α-helix to a β-sheet at the Kap3 N-terminus, may dictate kinesin-2 specificity for IFT.

Members of the kinesin-2 family coordinate with other motors to power diverse physiological processes^3^. Our cellular experiments show that β-hairpin-mediated autoinhibition is critical for the spatial regulation of Kif3 in IFT, as the motor accumulates at the ciliary tip when this mechanism is disabled and cannot be effectively returned to the cell body by dynein-2. We foresee that β-hairpin-mediated autoinhibition will be used widely in the kinesin-2 family to facilitate coordination with other motors, a hypothesis that can be tested using the β-hairpin mutations and structural framework for kinesin-2 regulation established here.

## METHODS

### Expression and purification of Kif3 and APC constructs

Genes encoding human Kif3A, Kif3B, Kif3C, Kap3, and APC^ARM^ were synthesized (Epoch, Eurofins or IDT) and inserted into the pACEBac1 vector (Geneva Biotech) using Gibson and HiFi assembly (NEB) with the following sequences added: Kif3A, C-terminal SNAP_f_ tag, TEV protease site and FLAG tag or C-terminal TEV protease site and FLAG tag; Kif3B/C, N-terminal ZZ tag and TEV protease site; Kap3 and APC^ARM^, N-terminal ZZ tag, TEV protease site and SNAP_f_ tag. The wild-type sequences were altered to generate deletions and mutations using Q5 site-directed mutagenesis reactions (NEB). All constructs were verified by DNA sequencing.

Constructs were expressed in *Spodoptera frugiperda* (Sf9) cells (Thermo Fisher Scientific) using the baculovirus system as previously described^64^. For Kif3AB and Kif3AC expression, 250 ml cultures were co-infected with Kif3A and Kif3B/C V_1_ viruses at 2%:0.5% Kif3A:Kif3B/C (v/v). For Kap3 and APC^ARM^ expression, cultures were infected with V1 virus at 1%. Proteins were purified and labeled with SNAP-Surface Alexa Fluor 647, SNAP-Cell TMR-Star or SNAP-Biotin (NEB) as described previously^65,66^.

### Size-exclusion chromatography and complex reconstitution

Proteins (5–10 μM in 100 μl) were analyzed using size-exclusion chromatography using either ÄKTAmicro or ÄKTA Go systems with a Superose 6 Increase 3.2/300 column (Cytiva) in Buffer A (50 mM Tris-HCl [pH 7.5], 150 mM K-acetate, 2 mM Mg-acetate, 5% Glycerol, 1 mM DTT, 0.2 mM Mg.ATP). For reconstitution of Kif3AB-Kap3, Kif3AB and Kap3 were incubated on ice at a 1:1 molar ratio for 15 min. For reconstitution of Kif3AB-Kap3:APC^ARM^, the pre-formed Kif3AB-Kap3 complex was incubated for 10 min with a two-fold molar excess of APC^ARM^ or APC^ARM-Mut^. Fractions of 50 μl were collected and analyzed by SDS-PAGE on 4–12% NuPAGE Bis-Tris gels (Thermo Fisher Scientific).

### Mass photometry

Kif3AB and Kap3 complexes were reconstituted as for size-exclusion chromatography experiments and then diluted to 1–100 nM using Buffer A (described above). Mass photometry measurements were recorded using the Refeyn OneMP instrument^67^ and analyzed using Discover software (Refeyn). NativeMark unstained protein standards (Thermo Fisher Scientific) were used to generate molecular mass calibration curves.

### Motility assays

Microtubules were polymerized from porcine tubulin (Cytoskeleton) as described^65^. For fluorescent visualization or surface attachment, 10% of HiLyte 488 tubulin or biotin tubulin (Cytoskeleton) were included in the polymerization mixture respectively.

Motility assays were carried out as described^65^ with the following modifications. For microtubule gliding and single-molecule assays with Kif3AB/C and Kap3, assays were carried out in Buffer B (50mM Tris-HCl [pH 8.0], 50 mM KCl, 2 mM MgCl_2_, 1 mM EGTA, 1 mM DTT, 20 μM taxol, with 1 mM Mg-ATP or AMPNP as indicated) supplemented with 1 mg/mL casein, 71 mM β-mercaptoethanol, 20 mM glucose, 300 μg/mL glucose oxidase and 60 μg/mL catalase. Final concentrations of Kif3AB/C were 0.15 – 2 nM. For Kap3 co-localization assays, Kap3 concentration was 1 nM. Motility assays including APC^ARM/ARM-Mut^ were carried out in Buffer C (80 mM PIPES [pH 6.9], 2 mM MgCl_2_, 1 mM EGTA, 1 mM DTT, 20 μM taxol, 1 mM Mg-ATP), with supplements as for Buffer B. Kif3AB and Kap3 were incubated on ice for 10 min prior to addition to the assay and, if included, APC^ARM/ARM-Mut^ was incubated for a further 5 min (final concentrations 40 nM Kif3, 1.25 μM APC^ARM/ARM-Mut^).

### TIRF microscopy

Fluorescently-labeled molecules were visualized on an Eclipse Ti-E inverted microscope with a CFI Apo TIRF 1.49 N.A. oil objective, Perfect Focus System, H-TIRF module, LU-N4 laser unit (Nikon) and a quad band filter set (Chroma). Images were recorded with 100 ms exposures on an iXon DU888 Ultra EMCCD camera (Andor), controlled with NIS-Elements AR Software (Nikon). The microscope was kept in a temperature-controlled environmental chamber (Okolab) operating at 25 °C for *in vitro* assays and 37 °C for live cell imaging, which was carried out as described^68^. Files were imported into FIJI^69^ (ImageJ, NIH) for analysis.

Velocities and durations of microtubule association were calculated from kymographs generated in FIJI as described^65,66^. For all single molecule measurements (run-length, velocity and landing rate) events longer than 4 consecutive pixels were included in the analysis. For measurement of Kap3 co-localisation, a circular region of interest was drawn around each Kif3A spot in the 640 nm channel and assessed for signal in the Kap3 561 nm channel. Graphing, curve fitting, and statistical analysis were performed in Prism9 and Prism10 (GraphPad).

### Electron microscopy

Reconstituted Kif3AB-Kap3 samples were diluted to 50–100 nM in Buffer A lacking glycerol, prepared for negative stain electron microscopy, and imaged as described^65^. Data were collected at a nominal magnification of 52,000×, giving a sampling of 2.09 Å/pixel. Subsequent image processing was carried out in Cryosparc^70^ unless stated otherwise. Particles were picked from micrographs using blob picker, followed by template picker once initial 2D classes were generated. Particles were extracted into 256-pixel boxes and subjected to multiple rounds of 2D classification from which a subset of well-resolved classes encompassing 3,032 individual particles was obtained. These classes were then compared to projections of the Kif3AB-Kap3 AF3 model. For this, the atomic model was converted to a density map in UCSF Chimera^71^ and low-pass filtered to 25 Å using EMAN2^72^. This volume was used to generate 3,032 projections in Cryosparc, using the simulate data job, and these were then classified into 50 2D classes. These classes were aligned to each data class average and scored by cross correlation to identify the best matching projection in SPIDER^73^.

Alphafold models were generated using Alphafold3^74^ and the ColabFold^75^ implementation of AlphaFold2^76^ and AlphaFold Multimer^77^. Visualization and structural alignment was carried out in UCSF Chimera and Chimera X^78^.

### Construct generation and cell biology experiments

For all cell biology experiments, mouse IMCD-3-FlpIn cells (gift from Peter K. Jackson, Stanford)^79^ were cultured in DMEM/F12 (Gibco) supplemented with 10% FBS, 100 U/ml penicillin-streptomycin. Cells were incubated in serum-free medium for 24 h to induce ciliogenesis.

For CRISPR-Cas9 genome editing in IMCD-3-FlpIn cells to generate a Kif3B knockout, guide RNA 5’-AAGCTCAGAATCAGTCCGGG-3’ targeting exon 2 of Kif3B was designed in Benchling and cloned into the pX330 Cas9 plasmid (gift from Feng Zhang; Addgene plasmid #42230)^80^. Transfection of Cas9 vector expressing Kif3B guide and validation of knockout were performed as described in detail previously^68^.

To stably express mScarlet-tagged Kif3B constructs in Kif3B knockout IMCD-3 cells, the Super PiggyBac transposon vector system (System Biosciences) was used. Cells in six-well plates were co-transfected with PiggyBac plasmid containing mScarlet-tagged gene of interest (Kif3B^WT^, Kif3B^ΔC156^ and Kif3B^KKK^) and a geneticin resistance marker (0.5 μg) and Super PiggyBac transposase expression vector (0.2 μg) using Lipofectamine 2000 (Thermo Fisher Scientific). Clones were selected using geneticin resistance (500 μg/ml) 2 d post transfection, cultured until confluent, and screened by live-cell TIRF microscopy and immunoblotting to confirm the expression of mScarlet labeled proteins. For continued culture, growth media contained 500 μg/ml geneticin.

### Immunoblotting

Cells were lysed in RIPA buffer containing 150 mM sodium chloride, 1.0% Triton X-100, 0.5% sodium deoxycholate, 0.1% SDS, 50 mM Tris [pH 8.0]. Samples were separated by SDS-PAGE followed by transfer to nitrocellulose membranes. Membranes were blocked overnight in 3% [w/v] Milk:TBS-T (20 mM Tris-base, 150 mM NaCl, 0.02% Tween20) and incubated for 4 h in ANTI-FLAG® M2 mouse antibody (1:1000; Sigma-Aldrich, F1804) or anti-GAPDH (1:10,000; Cell Signaling Technology, 2118) as described^64,68^. After incubation, membranes were washed for 4 × 5 min with TBS-T, before incubating with Goat Anti-Mouse IgG StarBright Blue 700 (1:1000; BioRad) or Goat Anti-Rabbit IgG (H + L) Alexa Fluor 647 (1:1000; Invitrogen) secondary antibodies in 2% [w/v] Milk:TBS-T for 1 h at room temperature. Blots were then washed with TBS-T for 4 × 5 min, before imaging.

### Immunofluorescence

IMCD-3 cells grown on 0.17 mm thick (#1.5) cover glass (VWR) were washed with PBS, followed by two washes in cytoskeletal buffer (100 mM NaCl, 300 mM sucrose, 3 mM MgCl_2_, 10 mM PIPES [pH 6.9]) and fixed in 4% paraformaldehyde prepared in cytoskeletal buffer with 0.5% Triton and 5 mM EGTA as described previously^68^. Cells were blocked in 3% BSA and 2% FBS in PBS for 1 h at room temperature and incubated overnight with respective antibodies. Post-primary antibody incubation, cells were washed and incubated with corresponding secondary antibody in a blocking solution for 1 h. Coverslips were mounted onto glass slides using mounting medium (GeneTex) and imaged by TIRF microscopy. The following antibodies were used at indicated dilutions: anti-acetylated tubulin (Sigma-Aldrich T6793; 1:2000); anti-gamma-tubulin (Sigma-Aldrich T6557; 1:500). AlexaFluor-labeled secondary antibodies (Thermo Fisher) were used at 1:500 dilution.

Immunofluorescence and live cell images were analyzed in FIJI. Measurement of cilia length and time-averaged fluorescence distributions of mScarlet Kif3B signal were carried out as described^64,68^.

## ACKNOWLEDGEMENTS

We thank Alexander Carver, Francisco Gonçalves-Santos, and William Allen (Sir William Dunn School of Pathology) for comments on the manuscript; Carylon Moores (Birkbeck) for discussions; Natasha Lukoyanova and Shu Chen (Birkbeck) for EM support; David Houldershaw, Yanni Goudetsidis, and Richard Westlake (Birkbeck) for computational support; and Cecilia Studniarek (Sir William Dunn School of Pathology) for assistance with Western blotting. This work is funded by grants from the Wellcome Trust (214998/Z/18/Z and 217186/Z/19/Z) and UKRI Biotechnology and Biological Sciences Research Council (BB/P008348/1).

## Notes

### Competing Interest Statement

The authors have declared no competing interest.

