## Supplementary_Information for "Beta-hairpin Mechanism of Autoinhibition and Activation in the Kinesin-2 Family"

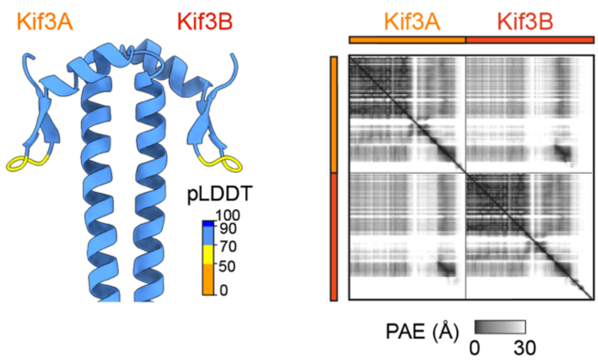


**Figure S1. Analysis of Kif3AB AF3 model.**
Left, Kif3AB AF3 model showing β-hairpin motif, colored by pLDDT (predicted local distance difference test) according to the key. Right, predicted aligned error (PAE) plot of Kif3AB, full-length proteins.

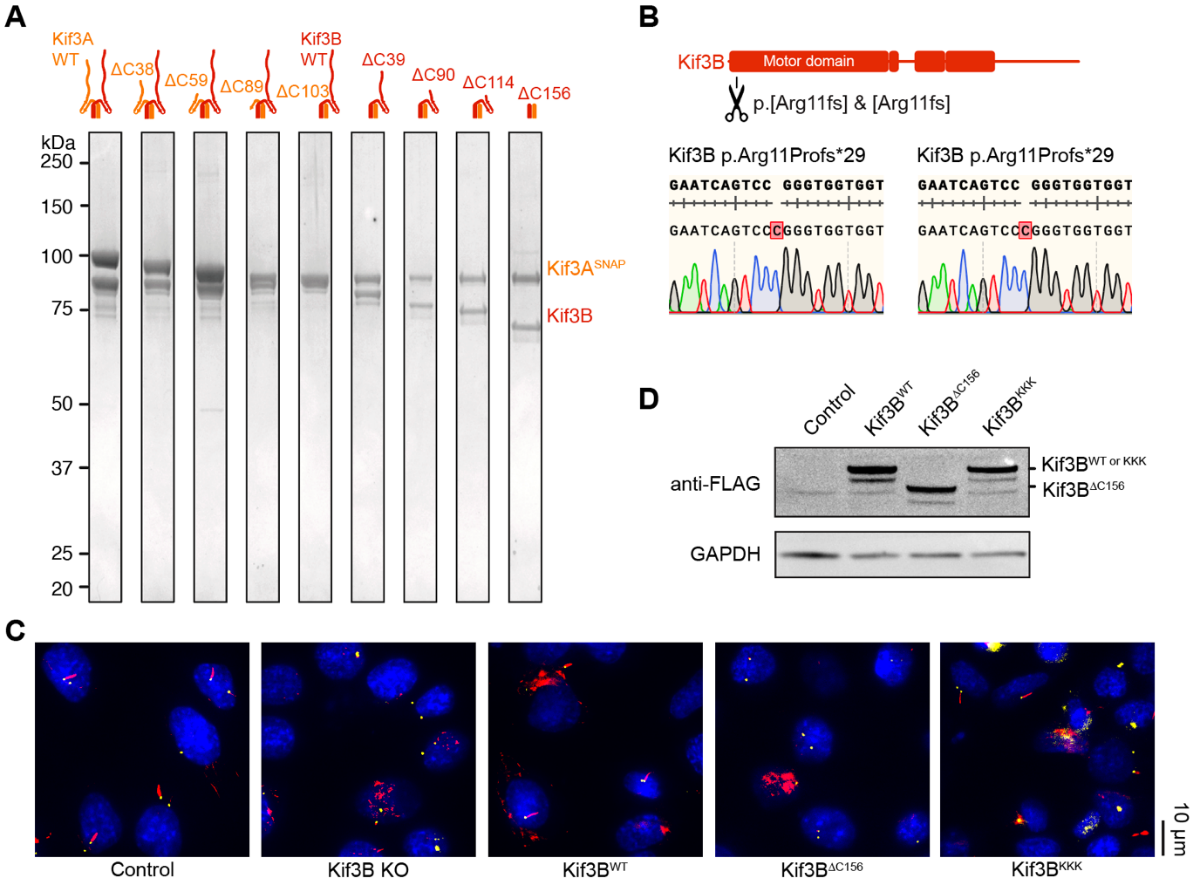


**Figure S2. Purification of Kif3AB constructs and CRISPR knockout of Kif3B.**
(A) SDS-PAGE of purified Kif3AB constructs (post size-exclusion chromatography) with indicated C-terminal truncations.
(B) Genotype for homozygous Kif3B KO cell line, with indels highlighted by alignment with the reference sequence. Clones were extensively Sanger sequenced to determine genotype (representative traces shown).
(C) Immunofluorescence images of cilia in Kif3B KO cell lines and cell lines stably expressing Kif3B$^{WT}$, Kif3B$^{\Delta C156}$ and Kif3B$^{KKK}$. Cells were stained for gamma tubulin (yellow), acetylated tubulin (red), and DAPI (blue).
(D) Western blot showing expression of FLAG-tagged Kif3B$^{WT}$, Kif3B$^{\Delta C156}$ or Kif3B$^{KKK}$ in Kif3B KO cells. Detected using anti-FLAG. GAPDH used as loading control.

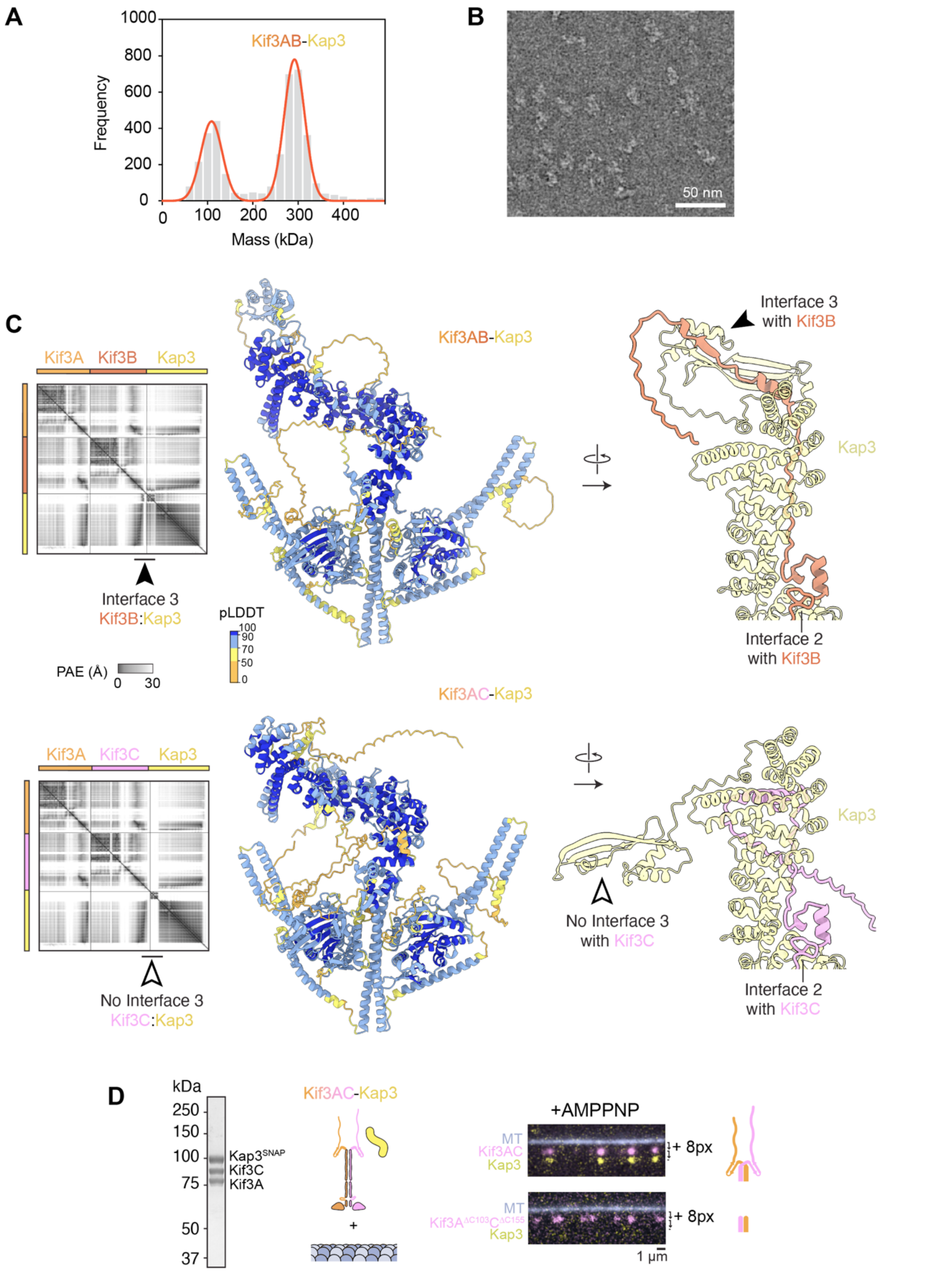


**Figure S3. Characterization of the Kif3AB-Kap3 and Kif3AC-Kap3 heterotrimer.**

(A) Mass photometry of purified Kif3AB-Kap3. Main peak (291 ± 23 kDa, mean ± SD) is consistent with the theoretical mass of heterotrimer (278 kDa).

(B) Electron micrograph of negatively stained Kif3AB-Kap3.

(C) Left, AF3 models of Kif3AB-Kap3 and Kif3AC-Kap3 colored by pLDDT according to the key. PAE plots alongside. Interface 3 highlighted with black arrowhead for Kif3B and lack of equivalent interface with white arrowhead for Kif3C. Right, Close-up views of Kap3 for each model. Note Interface 2 with Kap3 common to Kif3B and Kif3C and lack of Interface 3 with Kif3C (black vs. white arrowheads).

(D) Left, SDS-PAGE of purified Kif3AC-Kap3 heterotrimer following size-exclusion chromatography, peak fraction shown. Right, Kap3 co-localization assay with full-length Kif3AC (top) and with Kif3AC lacking C-terminal regions (Kif3A$^{\Delta C103}$C$^{\Delta C155}$; bottom). Composite TIRF images are shown of 488 nm channel (MT), 640 nm channel (Alexa-647-labeled Kif3A) offset in y-axis by 8 pixels, and 561 nm channel (TMR-labeled Kap3) offset in y-axis by 16-pixels.

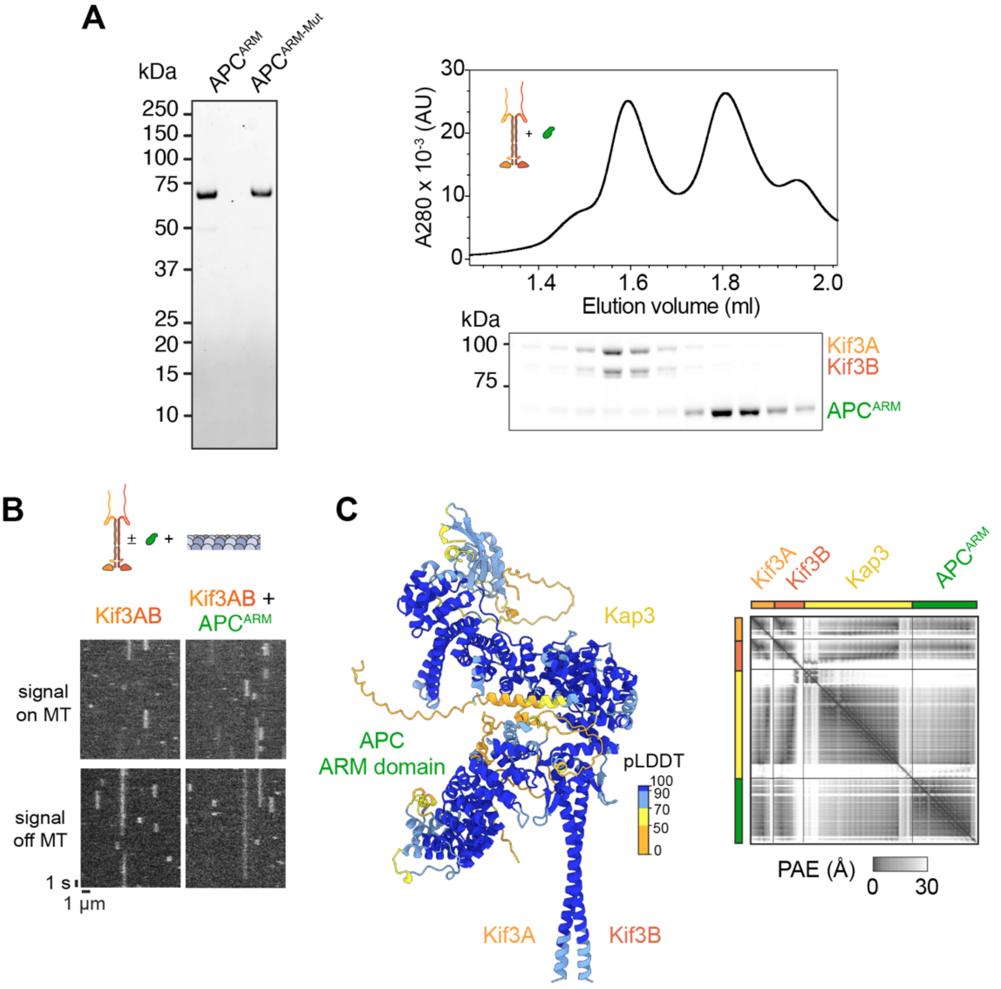


**Figure S4. Characterization of APC$^{ARM}$ and interaction with Kif3AB-Kap3.**

(A) Left, SDS-PAGE of purified APC$^{ARM}$ and APC$^{ARM\text{-}Mut}$ proteins. Right, size-exclusion chromatogram of binding reaction between Kif3AB and APC$^{ARM}$ with SDS-PAGE of fractions beneath. Note substantially reduced co-elution of APC$^{ARM}$ with Kif3AB compared to when Kap3 is present (Figure 6A).

(B) Examples kymographs of Kif3AB (Alexa-647-labeled Kif3A) in presence of microtubules and APC$^{ARM}$ (unlabeled). As there is non-specific binding of Kif3AB to the coverslip surface in this experiment, kymographs are shown for regions on and off the microtubule. APC$^{ARM}$ does not activate the motility of Kif3AB in the absence of Kap3.

(C) AF3 model of Kif3AB-Kap3-APC$^{ARM}$ colored by pLDDT according to the key. Right, PAE plot.

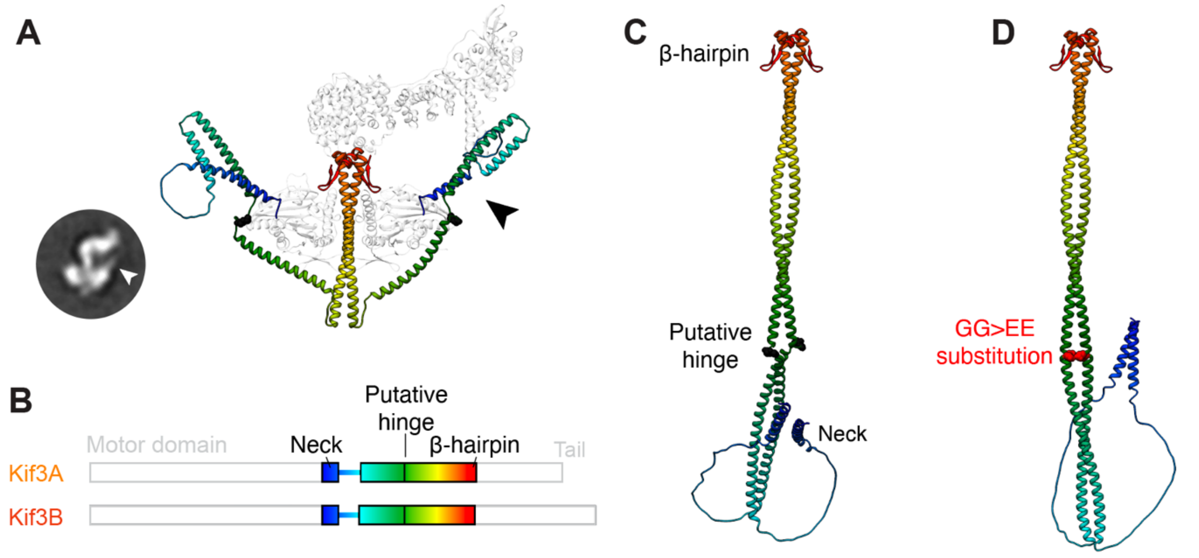


**Figure S5. Analysis of the coiled-coil regions in Kif3AB.**
(A) Kif3AB-Kap3 AF3 model with predicted coiled-coil regions in colored in rainbow from N-terminus (blue) to C-terminus (red). Note the predicted separation of the coiled coil strands in the N-terminal region in the autoinhibited state, resulting in a fine structure protruding from the motor domain (black arrowhead). Support for this model comes from EM class averages showing equivalent fine structure (white arrowhead, inset; from Figure 4D).
(B) Sequence diagrams of Kif3A and Kif3B, with predicted coiled coil regions colored in rainbow as in panel A. The location of the putative hinge in the coiled coil (Kif3A G485/G486; Kif3B G477/G478) is indicated.
(C) AF3 model of the Kif3AB coiled coil region lacking the motor domains (thereby preventing the formation of the autoinhibited conformation). This model shows an extended coiled coil, with a hinge at the predicted location, and may represent conformation of the coiled coil in active Kif3AB. Note that transition between the autoinhibited conformation in panel A and the extended conformation in panel C is not well described by simple folding at putative hinge.
(D) Mutation of the α-helix-breaking glycine residues at the putative hinge (Kif3A G485/G486; Kif3B G477/G478) to a residue with higher α-helical propensity (glutamic acid; E) is known to relieve autoinhibition in *C. elegans* orthologs Klp20-Klp11 (Brunnbauer *et al. PNAS* 107, 10460–10465, 2010). An AF3 model of the Kif3AB coiled coil region including the equivalent substitutions (Kif3A G485E/G486E; Kif3B G477E/G478E – GG>EE) shows a continuous coiled coil rather than a hinge, which could prevent access of the motor domains to the β-hairpin motif, consistent with the observed activation.
